# Poecivirus is present in individuals with beak deformities in seven species of North American birds

**DOI:** 10.1101/2020.02.03.927442

**Authors:** Maxine Zylberberg, Caroline Van Hemert, Colleen M. Handel, Rachel M. Liu, Joseph L. DeRisi

## Abstract

Avian keratin disorder (AKD), a disease characterized by debilitating beak overgrowth but with unknown etiology, has increasingly affected wild bird populations since the 1990s. We previously showed that a novel picornavirus, Poecivirus, is closely correlated with disease status in Black-capped Chickadees (*Poecile atricapillus*) in Alaska. However, our knowledge of the relationship between Poecivirus and beak deformities in other species and other geographic areas remains limited. The growing geographic scope and number of species affected by AKD-like beak deformities require a better understanding of the causative agent to evaluate the population-level impacts of this epizootic. Here, we tested eight individuals from six avian species with AKD-consistent deformities for the presence of Poecivirus: Mew Gull (*Larus canus*), Hairy Woodpecker (*Picoides villosus*), Black-billed Magpie (*Pica hudsonia*), Northwestern Crow (*Corvus caurinus*), Red-breasted Nuthatch (*Sitta canadensis*), and Blackpoll Warbler (*Setophaga striata*); individuals were sampled in Alaska and Maine (1999-2016). We used targeted PCR followed by Sanger sequencing to test for the presence of Poecivirus in each specimen, and to obtain viral genome sequence from virus-positive host individuals. We detected Poecivirus in all individuals tested, but not in negative controls. Furthermore, we used unbiased metagenomic sequencing to test for the presence of other pathogens in six of these specimens (Hairy Woodpecker, two Northwestern Crows, two Red-breasted Nuthatches, Blackpoll Warbler). This analysis yielded additional viral sequence from several specimens, including the complete coding region of Poecivirus from one Red-breasted Nuthatch, which we confirmed via targeted PCR followed by Sanger sequencing. This study demonstrates that Poecivirus is present in individuals with AKD-consistent deformities from six avian species other than Black-capped Chickadee. While further investigation will be required to explore whether there exists a causal link between this virus and AKD, this study demonstrates that Poecivirus is not geographically restricted to Alaska, but rather occurs elsewhere in North America.

## INTRODUCTION

Recently, a widespread epizootic of beak deformities consistent with avian keratin disorder (AKD) has been documented in dozens of avian species in North America and other parts of the world. Birds affected by AKD develop beak deformities characterized by elongation and often crossing and marked curvature (Handel et al. 2010). These deformities limit an individual’s ability to feed, preen, and cope with parasites, and result in decreased fitness and survival (Handel et al. 2010; Van Hemert et al. 2012, 2013; Wilkinson et al. 2016). While the population-level impacts of AKD remain unknown, the high prevalence, fitness impacts, and widespread occurrence of AKD across multiple species suggest that this pathology could have significant impacts on wild bird populations (Handel et al. 2010; Van Hemert and Handel 2010; Zylberberg et al. 2016).

AKD was first observed in Alaskan Black-capped Chickadees (*Poecile atricapillus*) in the late 1990s. In Black-capped Chickadees, AKD is typically seen in adult birds, with an average prevalence of 6.5% in south-central Alaska, where most research on this disease has been conducted (Handel et al. 2010). However, AKD-like deformities are not limited to Alaska or to Black-capped Chickadees. Morphologically similar deformities have been documented in more than 40 avian species in North America and over 30 species in the United Kingdom (Craves 1994; Handel et al. 2010; Harrison 2011; British Trust for Ornithology 2017), and appear to be particularly common in corvids, cavity-nesting passerines, and raptors (Handel et al. 2010; Van Hemert and Handel 2010; British Trust for Ornithology 2017). In Northwestern Crows (*Corvus caurinus*) in Alaska, AKD-like deformities are even more prevalent than in Black-capped Chickadees, with 17% of adults displaying clinical signs consistent with AKD (Van Hemert and Handel 2010). Moreover, this emergence of beak deformities has occurred concurrently across multiple species and the deformities appear to cluster geographically (Handel et al. 2010; British Trust for Ornithology 2017). While numerous species display similar gross pathology, the cause of these deformities is not known, and so it remains undetermined if a common factor is responsible. The increasing geographic scope and number of species potentially affected by AKD warrant a better understanding of the causative agent in order to evaluate the population-level impacts of this disease.

A variety of factors can contribute to beak deformities, including environmental contaminants, nutritional deficiencies, trauma, and exposure to infectious agents (Tully et al. 2000). However, multiple studies have failed to find clear evidence of a contaminant exposure, nutritional deficiency, or bacterial or fungal infection underlying AKD (Handel et al. 2010; Van Hemert et al. 2013; Handel and Van Hemert 2015). Using unbiased metagenomic sequencing of beak tissue from Black-capped Chickadees affected by AKD, we previously identified Poecivirus, a novel picornavirus, as a potential candidate agent (Zylberberg et al. 2016). Subsequent screening of 124 Black-capped Chickadees revealed that 28/28 individuals affected by AKD tested positive for Poecivirus, compared with only 9/96 asymptomatic individuals (Zylberberg et al. 2018); two Northwestern Crows and two Red-breasted Nuthatches (*Sitta canadensis*) with AKD-consistent pathology also tested positive for Poecivirus (Zylberberg et al. 2016). Furthermore, we previously used both *in situ* hybridization and a strand-specific expression assay to localize Poecivirus to beak tissue of AKD-positive individuals and to provide evidence of active viral replication (Zylberberg et al. 2018). This evidence establishes a strong correlation between disease status in Black-capped Chickadees and the presence of Poecivirus (Zylberberg et al. 2016, 2018) although a definitive causal link remains to be tested.

Here, we use targeted PCR followed by Sanger sequencing to test for the presence of Poecivirus in the beak and cloacal tissue of four opportunistically collected individuals with deformed beaks, representing four species: Mew Gull (*Larus canus*), Hairy Woodpecker (*Picoides villosus*), Black-billed Magpie (*Pica hudsonia*), and Blackpoll Warbler (*Setophaga striata*). We then used unbiased metagenomic sequencing to analyze tissues of six individuals: the Hairy Woodpecker and Blackpoll Warbler, plus two Northwestern Crows and two Red-breasted Nuthatches from Alaska that we had previously found to be positive for Poecivirus (Zylberberg et al. 2016). We tested these six specimens for the presence of other pathogens and to obtain additional viral genome sequence.

## MATERIALS AND METHODS

### Specimen collection

We opportunistically obtained one Mew Gull (Charadriiformes: Laridae), one Hairy Woodpecker (Piciformes: Picidae), one Black-billed Magpie (Passeriformes: Corvidae), two Northwestern Crows (Passeriformes: Corvidae), two Red-breasted Nuthatches (Passeriformes: Sittidae), and one Blackpoll Warbler (Passeriformes: Parulidae) with grossly overgrown beaks between 1999 and 2016 (Table 1, Figure 1). The Mew Gull, Hairy Woodpecker, and Black-billed Magpie were submitted to a wildlife rehabilitation center in Anchorage, Alaska, and later euthanized. The Blackpoll Warbler was submitted to a wildlife rehabilitation center in Maine and later euthanized. One Northwestern Crow was submitted to a wildlife rehabilitation center in Juneau, Alaska, and later euthanized; the second crow was collected by shotgun as part of airport safety measures in Juneau, Alaska. The two Red-breasted Nuthatches were collected after dying in the wild in Anchorage and Chugiak, Alaska. Specimens were stored frozen at −20°C until processed.

**Table 1.** Avian specimens analyzed in this study. Common and scientific names of birds, specimen identification (ID), collection location and date, number of next-generation sequencing (NGS) paired-end reads, and tissues in which Poecivirus was detected.

| Species | Specimen ID | Location | City, State | Date | NGS reads | Poecivirus detected |  |
| --- | --- | --- | --- | --- | --- | --- | --- |
|  |  |  |  |  |  | Beak | Cloaca |
| Mew Gull (*Larus canus*) | MEGU 993 | 61.17 N, 149.86 W | Anchorage, Alaska | 5 Sep 2016 | — | Yes | Yes |
| Hairy Woodpecker (*Picoides villosus*) | HAWO 969 | 61.09 N, 149.78 W | Anchorage, Alaska | 8 Apr 2015 | $7.1 \times 10^7$ | Yes | Yes |
| Black-billed Magpie (*Pica hudsonia*) | BBMA 992 | 61.33 N, 149.56 W | Eagle River, Alaska | 26 Aug 2016 | — | No | Yes |
| Northwestern Crow (*Corvus caurinus*) | NOCR 674 | 58.28 N, 134.41 W | Juneau, Alaska | 11 Jul 2005 | $1.4 \times 10^7$ | Yes | Yes |
| Northwestern Crow (*Corvus caurinus*) | NOCR 848 | 58.36 N, 134.58 W | Juneau, Alaska | 27 Feb 2007 | $5.5 \times 10^7$ | Yes | Yes |
| Red-breasted Nuthatch (*Sitta canadensis*) | RBNU 12 | 61.13 N, 149.91 W | Anchorage, Alaska | 29 Mar 1999 | $7.2 \times 10^7$ | No | Yes |
| Red-breasted Nuthatch (*Sitta canadensis*) | RBNU 929 | 61.41 N, 149.48 W | Chugiak, Alaska | 10 Dec 2012 | $5.9 \times 10^7$ | Yes | Yes |
| Blackpoll Warbler (*Setophaga striata*) | BLPW 1 | 43.17 N, 70.61 W | Cape Neddick, Maine | 8 Nov 2016 | $6.0 \times 10^7$ | Yes | No |

**Figure 1.**
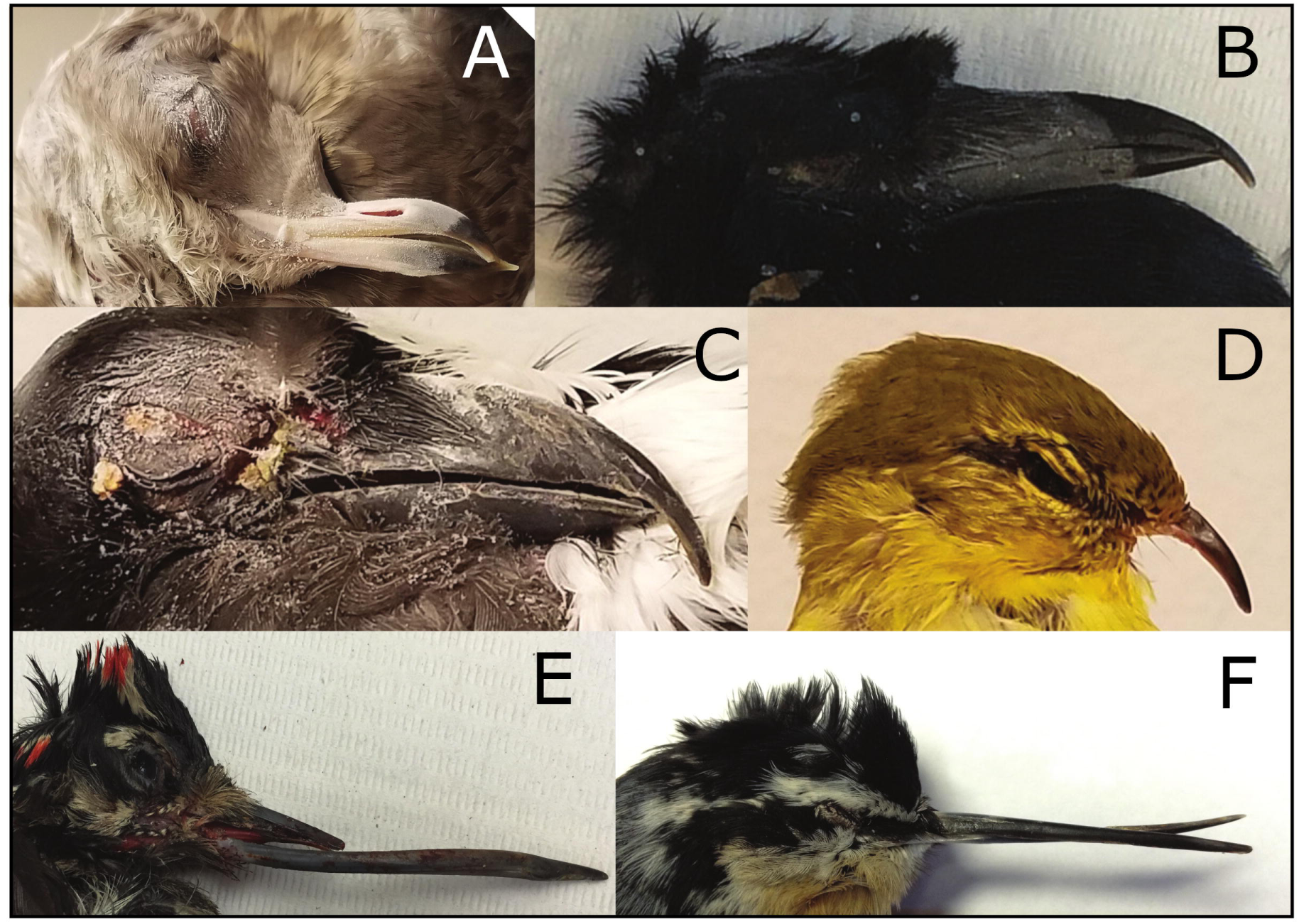
Specimens analyzed in this study exhibiting beak overgrowth characteristic of avian keratin disorder: (A) Mew Gull 993 (*Larus canus*), (B) Northwestern Crow 674 (*Corvus caurinus*), (C) Black-billed Magpie 992 (*Pica hudsonia*), (D) Blackpoll Warbler 1 (*Setophaga striata*), (E) Hairy Woodpecker 969 (*Picoides villosus*), and (F) Red-breasted Nuthatch 929 (*Sitta canadensis*). Images are not to scale.

All work was conducted with the approval of the U.S. Geological Survey’s (USGS) Alaska Science Center Institutional Animal Care and Use Committee (assurance no. 2016-14) and under appropriate state and federal permits.

### Detection of Poecivirus

We tested for the presence of Poecivirus in beak and cloacal tissue from all individuals using targeted PCR primers followed by Sanger sequencing. RNA was extracted from tissue samples using a Direct-zol RNA MicroPrep kit (Zymo Research, Irvine, CA, USA); briefly, 5 mg of tissue was ground using a pellet pestle (cloacal tissue) or a mortar and pestle on liquid nitrogen (beak tissue) and the extraction proceeded as described in the manufacturer’s protocol. Next, 200 ng of RNA were reverse transcribed in 10 μl reactions containing 100 pmol random hexamer, 1x reaction buffer, 5 mM dithiothreitol, 1.25 mM (each) deoxynucleoside triphosphates (dNTPs), and 100 U Superscript III (Life Technologies, Carlsbad, CA, USA); mixtures were incubated at 25°C for 5 min, 42°C for 60 min, and 70°C for 15 min. Samples were then screened via PCR using Poecivirus-specific primers (Table S1). PCR mixtures contained 1x iProof master mix (Bio-Rad Laboratories, Hercules, CA, USA), 0.5 M primer, and 5µl cDNA. Thermocycler conditions were set at 98°C for 30s, then 40 cycles of 98°C for 10 s, 58-62°C for 10 s, and 72°C for 30 s, then a 5-min elongation step of 72°C. Annealing temperature was optimized for each set of primers using a gradient cycler. Amplicons were visualized on 1.5% agarose gels; those in the correct size range were extracted and purified using a Zymoclean gel DNA recovery kit (Zymo Research, Irvine, CA, USA) and Sanger sequenced (Quintara Biosciences, South San Francisco, CA, USA). Positive controls included beak tissue from three Black-capped Chickadees that had previously tested positive for the presence of Poecivirus (Zylberberg et al. 2016). Negative controls included water as well as separate beak and cloacal tissue samples from four Black-capped Chickadees without gross beak deformities and that had previously tested negative for the presence of Poecivirus (Zylberberg et al. 2016). When testing of beak and cloaca tissue samples from the same individual resulted in contradictory results, tests were confirmed in triplicate; in each case, the results for each individual tissue were consistent across replicates.

### Virus discovery: sample processing for high-throughput sequencing

A sample of beak tissue and a sample of cloacal tissue from each individual Hairy Woodpecker, Northwestern Crow, Red-breasted Nuthatch, and Blackpoll Warbler were subjected to metagenomic deep sequencing (Table 1). RNA extraction proceeded as described above. Samples were prepped for high-throughput sequencing using the NEBNext Ultra RNA Library Prep Kit for Illumina (New England Biolabs, Ipswich, MA, USA) following the manufacturer’s protocol. Sequencing was conducted on the Illumina HiSeq 4000 using a single sequencing lane and resulted in 135-nucleotide-long paired-end reads.

### Sequence analysis

Sequences were analyzed using Bowtie 2 (Langmead and Salzberg 2012) to align reads to the Poecivirus reference genome (GenBank KU977108) (Zylberberg et al. 2016). Aligned reads were assembled to the Poecivirus reference genome to obtain consensus sequence from Poecivirus present in each specimen. In addition, to conduct an unbiased search for other pathogens that may be present in the specimens, sequences were analyzed using a rapid computational pipeline developed to identify potential pathogens present in metagenomic data as previously described (Doan et al. 2016). The pipeline filters out low quality reads, identifies the best match for each read in the NCBI nucleotide database, then indicates credible organism “hits” warranting further confirmatory testing if >20 non-redundant read pairs per million read pairs (rM) map to an organism at the species level based on nucleotide alignment. Briefly, reads were aligned to available avian reference genomes: *Corvus brachyrhynchos* (assembly ASM69197v1), *Zonotrichia albicollis* (assembly Zonotrichia_albicollis-1.0.1), and *Geospiza fortis* (assembly GeoFor_1.0). Unaligned (i.e., non-avian) reads were quality-filtered using PriceSeqFilter (Ruby et al. 2013) and compressed by cd-hit-dup (v4.6.1); then low-complexity reads were removed via the Lempel-Ziv-Welch algorithm (Ziv and Lempel 1977; Fu et al. 2012). GSNAPL (v2015-12-31) (Wu and Nacu 2010) was used to align the remaining non-avian reads to the NCBI nucleotide (nt) database. The same reads were also aligned to the NCBI nonredundant protein (nr) database using the Rapsearch2 algorithm (Zhao et al. 2011).

## RESULTS

Using targeted PCR followed by Sanger sequencing, we detected Poecivirus in 8/8 specimens that displayed AKD-consistent beak deformities, representing six avian species (Table 1). Poecivirus was not detected in negative controls (neither in water nor in beak and cloacal tissue from Black-capped Chickadees without beak deformities and previously confirmed Poecivirus-negative). In 6/8 of the specimens with beak deformities, we detected virus in the beak tissue, while in 7/8 we detected virus in the cloaca; the Blackpoll Warbler was the only specimen in which we detected virus in the beak but not in the cloaca. All sequenced regions had high pairwise nucleotide identity (93-96.5%; Figure 2) to Black-capped Chickadee Poecivirus (GenBank KU977108).

**Figure 2.**
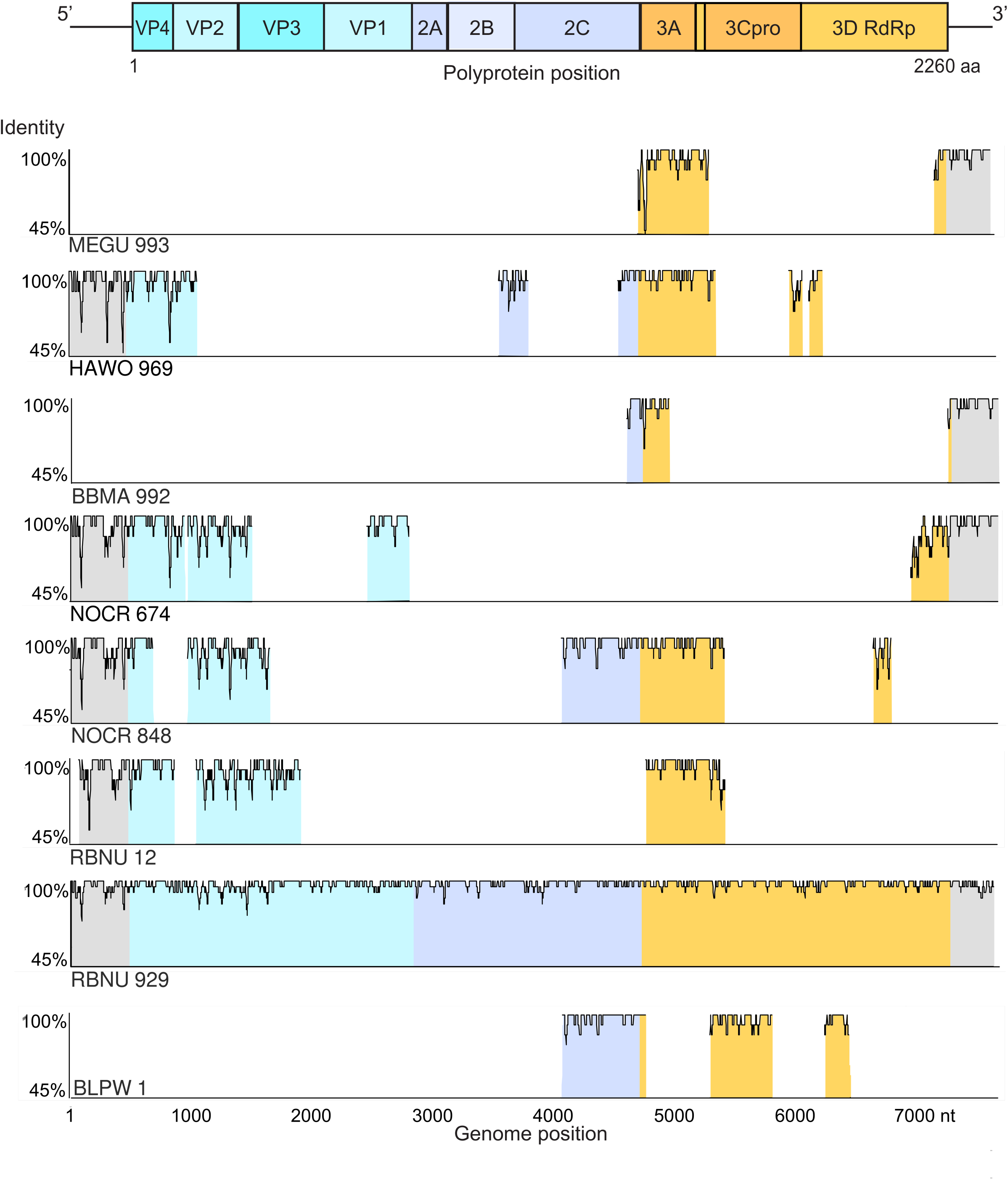
Homology to Poecivirus. (Top) Predicted Poecivirus genome organization. Remaining rows show nucleotide homology between viral sequence obtained from each specimen and Poecivirus, measured as the pairwise identity of a moving 15-nucleotide window. Blue (P1) represents viral structural proteins, violet (P2) and orange (P3) represent nonstructural proteins, and grey represents the 5’ and 3’ non-coding regions.

Unbiased sequencing of RNA from six individuals with beak deformities on the Illumina HiSeq platform generated approximately 1.4 × 10^7^ to 7.2 × 10^7^ 126-nucleotide single-read sequences per specimen (Table 1). Retrospective mapping of the sequenced reads to the Poecivirus genome revealed the full polyprotein coding region (7588 nt) of Poecivirus in one Red-breasted Nuthatch (929); in addition, it revealed portions of the Poecivirus polyprotein coding region of variable size (789-3166 nt) in the other specimens tested (Figure 2). Metagenomic sequencing combined with our bioinformatic pathogen discovery pipeline failed to detect a pathogen other than Poecivirus present in the specimens tested.

Comparison to the NCBI nucleotide (nt) and protein (nr) databases revealed that the full polyprotein sequence from Red-breasted Nuthatch 929 (validated by Sanger sequencing) was more closely related to Poecivirus than to any other known virus, exhibiting 96.5% pairwise nucleotide identity and 98.2% amino acid identity with Poecivirus isolated from Alaskan Black-capped Chickadees (Zylberberg et al. 2016). The 7588 nt viral genome recovered from the Red-breasted Nuthatch exhibited a total of 267 mutations compared to the Black-capped Chickadee Poecivirus genome as determined by Sanger sequencing. Of these mutations, 202 (75.7%) were supported by Illumina sequencing data with an average depth of coverage of 427 reads per nucleotide, and 112 were non-synonymous. Both our negative controls and the large number of Illumina and Sanger confirmed variants observed in this full length polyprotein supports the notion that this sequence did not arise from contaminating Poeciviruses previously amplified in this laboratory.

All raw non-genomic data that support the findings of this publication can be found in the accompanying USGS data release (Zylberberg et al. 2019). Genomic data are available at NCBI (GenBank accession numbers MN944596-MN944619).

## DISCUSSION

In this study, we investigated whether Poecivirus was present in individuals of multiple species that exhibited clinical signs consistent with avian keratin disorder (AKD), a disease characterized by gross beak deformities. The concurrent emergence of beak deformities across species and the geographic proximity of such clusters lend support to the idea that multiple species are being afflicted by a disease with a common etiology (Handel et al. 2010). We previously identified and characterized Poecivirus, a novel picornavirus, present in Black-capped Chickadees with AKD (Zylberberg et al. 2016), demonstrated a strong correlation between the presence of Poecivirus and AKD, and showed that Poecivirus was infecting, and appeared to be replicating in, the beak tissue of Black-capped Chickadees with AKD (Zylberberg et al. 2018). While our previous work demonstrated a strong association between Poecivirus and AKD in Black-capped Chickadees, our current study provides evidence that the same etiological agent could be responsible for similar beak deformities in other avian species.

Here, we expanded on our previous work by testing eight individuals with signs consistent with AKD, representing six avian species (Mew Gull, Hairy Woodpecker, Black-billed Magpie, Northwestern Crow, Red-breasted Nuthatch, Blackpoll Warbler) for the presence of Poecivirus. In addition, we applied unbiased, metagenomic next-generation sequencing to search for other pathogens in six of these individuals. In keeping with the hypothesis that the emergence of beak deformities across species is causally linked, we found Poecivirus to be present in 8/8 (100%) of individuals tested using targeted PCR followed by Sanger sequencing, albeit not consistently across sample types. Meanwhile, unbiased metagenomic sequencing of beak and cloacal tissue failed to detect another candidate pathogen. While we cannot rule out the presence of additional pathogens in other tissues, this result is consistent with our previous study of Black-capped Chickadees, in which unbiased metagenomic sequencing identified Poecivirus but no other pathogens in the beaks of AKD-affected individuals.

We recovered the complete polyprotein coding region of virus from a single specimen, Red-breasted Nuthatch 929; in addition, we obtained more than 25% of the viral genome from the Hairy Woodpecker and Northwestern Crow 848 (2.5 and 3.2 Kb, respectively). Despite efforts to obtain the full polyprotein coding regions of the virus from other specimens, we were unable to do so; this is likely due to the variable condition of the specimens, which were collected opportunistically and underwent variable handling conditions prior to sample processing at the University of California. Sequence analysis revealed that viruses from the Hairy Woodpecker, Red-breasted Nuthatch 929, and Northwestern Crow 848 (GenBank accession numbers MN944614-MN944618, MN944596, and MN944600-MN944602, MN 944604, respectively) are closely related to the virus sampled from Black-capped Chickadees, exhibiting 96%, 94%, and 94% nucleotide identity, respectively, with the Poecivirus reference genome (GenBank accession number KU977108); in the case of the Red-breasted Nuthatch and Northwestern Crow, these data expand upon, and are consistent with, previous analysis of more limited viral sequence from these specimens (Zylberberg et al. 2016). Except for the red-breasted nuthatch, next-generation sequencing of better-preserved specimens will be necessary to obtain the full viral genome from each of these hosts to confirm that they are infected by Poecivirus, as opposed to a related but distinct species of virus.

Previous studies of Poecivirus have largely focused on Black-capped Chickadees from Alaska. Therefore, while beak deformities characteristic of AKD are geographically widespread, it remained unknown whether Poecivirus was limited to Alaska, or had a broader geographic range. The Blackpoll Warbler included here was the first specimen tested for Poecivirus that was collected outside of Alaska (Maine); detection of Poecivirus in this specimen provides evidence that this pathogen occurs elsewhere in North America.

The focus of the current study was to determine whether Poecivirus was present in individuals with beak deformities of avian species other than the Black-capped Chickadee. Specimens were obtained opportunistically; as a result, our sample sizes were small, and our conclusions are necessarily limited. Of particular importance is the lack of corresponding control individuals from the additional species because we had none available to test at the time of this study. However, the lack of detection of Poecivirus in negative controls from Black-capped Chickadees indicates that positive detections in the newly tested species were not the result of lab contamination. Moreover, the detection of 202 nucleotide differences between the strain of Poecivirus obtained from the Red-breasted Nuthatch, supported by both Sanger and next-generation sequencing data, further corroborates this as a distinct viral specimen; 112 of these mutations were non-synonymous changes. While we have demonstrated the presence of Poecivirus in additional species, families, and orders of birds, screening of larger numbers of individuals of each species, including individuals with no clinical signs of AKD, will be required to test the association between Poecivirus and beak deformities in these species. The use of cloacal and buccal swabs to test individuals for infection provides an effective non-terminal method of sampling Black-capped Chickadees for Poecivirus (Zylberberg et al. 2018); however, the efficacy of these methods needs to be evaluated in additional species.

In conclusion, Poecivirus continues to warrant further investigation as a candidate agent of AKD across host species. The data presented here demonstrate that Poecivirus is not limited to Black-capped Chickadees, but rather is present in individuals of multiple species exhibiting AKD-consistent deformities, including passerines and non-passerines sampled from both the western and eastern United States. However, a larger sample size will be needed to evaluate the relationship between Poecivirus and beak deformities in the Mew Gull, Hairy Woodpecker, Black-billed Magpie, Northwestern Crow, Red-breasted Nuthatch, and Blackpoll Warbler. Ultimately, a viral challenge of healthy individuals with Poecivirus is needed to determine with certainty the role of Poecivirus in AKD. Such knowledge will improve our understanding of the potential impacts of this disease on avian populations worldwide.

## ACKNOWLEDGMENTS

We thank Lisa Pajot, Rachel Richardson, Diana Dumais, Sonja Ahlberg, the Bird Treatment and Learning Center, USDA-APHIS Wildlife Services, and members of the public for assisting with specimen collection. Sarah Sonsthagen and James Angus Chandler provided helpful comments on earlier versions of this manuscript. Funding was provided by a National Science Foundation postdoctoral research fellowship (M.Z.), the Howard Hughes Medical Institute (J.L.D.), the Chan Zuckerberg Biohub (J.L.D.), and the USGS Ecosystems Mission Area (C.M.H and C.V.H.). Any use of trade, firm, or product names is for descriptive purposes only and does not imply endorsement by the U.S. government.

---

**Table S1**. Primer sequences.

