## Supplemental Table 1 for "Poecivirus is present in individuals with beak deformities in seven species of North American birds"

| **Primer pair** | **Primer name** | **Sequence** |
| --- | --- | --- |
| 1 | BCCHpic_2F | CCAACTGAGTCGGATTCTACTG |
|  | BCCHpic_2R | GTGAAACACCACCCAACATCA |
| 2 | BCCHpic_9F | GCTGGAAAGACTCGCGTAAT |
|  | BCCHpic_9R | TTGCTCCCCACCCATTGTAT |
| 3 | Poeci_1F | TGGCTGCTCTAGAGGATAAAGG |
|  | Poeci_1R | ACTGCACTACAACCAAATCTGT |
| 4 | Poeci_2F | AGCTTGGCCCCTCTAATTGT |
|  | Poeci_2R | GATTACTGTTCCGGTCTCTTGG |
| 5 | Poeci_4F | TGGGCATTGTCTCGAGTGTA |
|  | Poeci_4R | TACGAAAAGCCTCAGTCGGA |
| 6 | Poeci_8F | TCTTGGTTGTGGTGGAGACT |
|  | Poeci_8R | GTGAAACACCACCCAACATCA |
| 7 | Poeci_9F | TGTTGATGTTGAGCAGTGTAGC |
|  | Poeci_9R | TTGCTCCCCACCCATTGTAT |
| 8 | Poeci_10F | AGCCAATGTATCAAAGTCCAGTG |
|  | Poeci_10R | CGACCGCATATCTTGTACGT |
| 9 | Poeci_11F | TGTAATTTGGCAGGGTGTTCT |
|  | Poeci_11R | ACTCCTTCAAAAGCATCACCC |
| 10 | Poeci_12F | GATGTTTCGTTTGTGGGACCT |
|  | Poeci_12R | TGCCTGGACAGATAAGGAGG |
| 11 | Poeci_13F | GGGCTCAGATTTTCAATTAAGGG |
|  | Poeci_13R | CTATTCACCTGCACCGACAC |
| 12 | Poeci_14F | GAAATCTGGGGCCCACAATG |
|  | Poeci_14R | AAGCCTCAGTCGGATTTAACA |

Supplemental Table 1: primers.
